# YOUNG PLASMA REJUVENATES BLOOD DNA METHYLATION PROFILE, PROLONGS MEAN LIFESPAN AND IMPROVES HEALTH IN OLD RATS

**DOI:** 10.1101/2022.12.01.518747

**Authors:** Priscila Chiavellini, Martina Canatelli Mallat, Marianne Lehmann, Joseph A. Zoller, Juozas Gordevicius, Maria D. Gallardo, Diana C. Pasquini, Ezequiel Lacunza, Claudia B. Herenu, Gustavo R. Morel, Steve Horvath, Rodolfo G. Goya

## Abstract

There is converging evidence that young blood conveys cells, vesicles and molecules able to revitalize function and restore organ integrity in old individuals. Here, we assessed the effects of young rat plasma on the lifespan, epigenetic age and healthspan of old female rats. Beginning at 25.3 months of age, a group of 9 rats (group T) was intraperitoneally injected with plasma from young rats (2 months) until their natural death. A group of control rats of the same age, received no treatment. Blood samples were collected every other week. Survival curves showed that from age 26 to 30 months, none of the T animals died, whereas the survival curve of C rats began to decline at age 26 months. The external appearance of the T rats was healthier than that of the C counterparts. Blood DNA methylation (DNAm) was assessed using the HorvathMammalMethylChip320. Blood DNAm age versus chronological age showed that DNAm age in young animals increased faster than chronological age then slowed down progressively, entering a plateau after 27 months. Immediately after the start of the treatment, the DNAm age (i.e., epigenetic age) of the treated rats fell below the DNAm age of controls and remained consistently lower until the end of their lives. Assessment of each experimental group showed that the blood DNA methylation levels of 1638 CpGs were different between treated and control blood samples (false discovery rate q-value<0.05). Of these, 1007 CpGs exhibited increased methylation, with age while 631 CpGs showed decreased methylation levels. When rats were grouped according to the similarities in their differential blood DNA methylation profile, samples from the treated and control rats clustered in separate groups. Analysis of promoter differential methylation in genes involved in systemic regulatory activities revealed specific GO term enrichment related to the insulin-like factors (IGFs) pathways as well as to cytokines and chemokines associated with immune and homeostatic functions. We conclude that young plasma therapy may constitute a natural noninvasive intervention for epigenetic rejuvenation and health enhancement, readily translatable to the clinic.

## INTRODUCTION

There is converging evidence that young blood conveys cells, vesicles and molecules able to revitalize function and restore organ integrity in old individuals. This area of research was started in the 1860s by the French physiologist Paul Bert **(1)**, who introduced a surgical technique, later termed parabiosis, to connect the circulatory systems of two living albino rats. The first to apply parabiosis of a young and old rat, (known as heterochronic parabiosis) to the study of aging was Clive McCay who reported that when old and young rats were joined for 9–18 months, the older animals’ bones became similar in weight and density to the bones of their younger counterparts **(2).** The same group studied the lifespans of old–young rat pairs. Older partners lived for four to five months longer than controls, suggesting for the first time that circulation of young blood might increase longevity **(3).** After lying dormant for nearly 30 years **(1**972 to 2001), parabiosis started to be used again, now, for the study of the nature and migration of hematopoietic stem cells in vivo **(4, 5)**.

Emerging evidence that age-related decline of progenitor cell activity may be reversed by young blood factors came from a report that heterochronic parabiosis in mice restored the activation of Notch signaling as well as the proliferation and regenerative capacity of muscle satellite cells in old mice and that the intervention also increased proliferation in aged hepatocytes **(6).**

To our knowledge, the first report that young plasma has restorative effects in old animals was published by Villeda et al., in 2014 **(7)**. In effect, the study shows that administration of young plasma improves hippocampal-dependent learning and memory in aged mice. It has been hypothesized that young plasma may possess life-extending properties, but reported data are scant and controversial. In mice, it has been reported that repeated injections of young plasma do not significantly impact either median or maximal lifespan **(8).** In contrast, a more recent study in old rats reported that a porcine plasma fraction (termed elixir or E5) markedly reversed the epigenetic age of the animals which also showed functional and metabolic signs of rejuvenation **(9).** The latter report prompted us to assess the effects of young rat plasma on the lifespan, epigenetic age and healthspan of old female rats. Here, we report the results of the study.

## METHODS

### Animals

Seventeen female Sprague Dawley (SD) rats weighing (X±SEM) 250±9 g, were used. At the start of the young plasma treatment, all rats were 25.3 months old. Animals were housed in a temperature-controlled room (22 ± 2°C) on a 12:12 h light/dark cycle. Food and water were available ad libitum. All experiments with animals were performed in accordance with the Animal Welfare Guidelines of NIH (INIBIOLP’s Animal Welfare Assurance No A5647-01). The ethical acceptability of the animal protocols used here have been approved by our institutional IACUC (Protocol # T09-01-2013).

#### Young plasma preparation

Two-month old rats were i.p. injected with 2,500 IU sodium heparin 15 min before decapitation. Trunk blood was collected in Flacon tubes containing 150 μl Na heparin at 7,500 IU /ml. The blood is centrifuged at 3,000 g for 15 minutes, plasma collected and stored in 10-ml aliquots and stored at −80 °C.

#### Experimental design

Animals were allotted to two groups: 1) Control group (C, 8 rats), received no treatment. 2) Beginning at 25.3 months of age, each rat of the treated group (T, 9 rats), received a slow (2 min) intraperitoneal (i.p.) injection of young plasma (1 ml) every other week. The treatment was life-long (until the natural death of all animals).

#### Blood withdrawal and processing

Blood samples were taken from the tail veins every other week in order to determine epigenetic age during the experiment. Blood samples were stored at −80 °C until the end of the experiment.

### Purification and methylation analysis of genomic DNA

DNAs from 187 blood samples were purified using the DNeasy Blood and Tissue Kit (Qiagen). Only DNA samples with 260/280 ratios greater than 2.0 were processed for methylation analysis. The bisulfite conversion of genomic DNA was performed using the EZ Methylation Kit (Zymo Research, D5002), following the manufacturer’s instructions. The converted genomic DNA was analyzed by Horvath Mammal Methyl Chip 320. The Horvath Mammal Methyl Chip 320 assay provides quantitative measurements of DNA methylation for 105615 CpG dinucleotides that map to the Rattus norvegicus UCSC 6.0 genome (and other mammalian sequences**) (10).**

### Statistical analysis of differentially methylated positions

Quality control (QC), pre-processing, and statistical analysis of methylation profiles were performed with “minfi” R/Bioconductor package. **(11)** Briefly, QC analysis was performed with getQC function and further preprocessed with the Noob/ssNoob method.

Data exploratory analysis was performed through an unsupervised hierarchical clustering analysis based on Euclidean distance of methylation profiles. In addition, the distribution of the methylation measurements in each sample was evaluated. To test differential methylation levels in each CpG, multiple hypothesis testing was performed through a multivariate linear regression model using the “limma” package **(12).** The positive false discovery rate was controlled with a q-value threshold of 0.05 **(13).**

Gene ontology (GO) enrichment test for positively and negatively differentially methylated CpGs from the Horvath Mammal 320 array was conducted with GOmeth function of “missMethyl” package **(14)**. GO gene sets were evaluated with a threshold of FDR < 0.05. A functional enrichment analysis was also performed for hyper and hypomethylated CpGs at promoter regions of blood DNA from control and treated rats **(Suppl Table 5) (15).**

## RESULTS

### Blood DNAm age curves flatten at extreme ages

In young animals, DNAm age moves faster than physical time whereas in very old rats

DNAm age enters a virtual plateau **(Fig. 1)**. In epigenetic age terms, old rats do not age regardless of whether they belong to the control or experimental groups.

**Figure 1.**
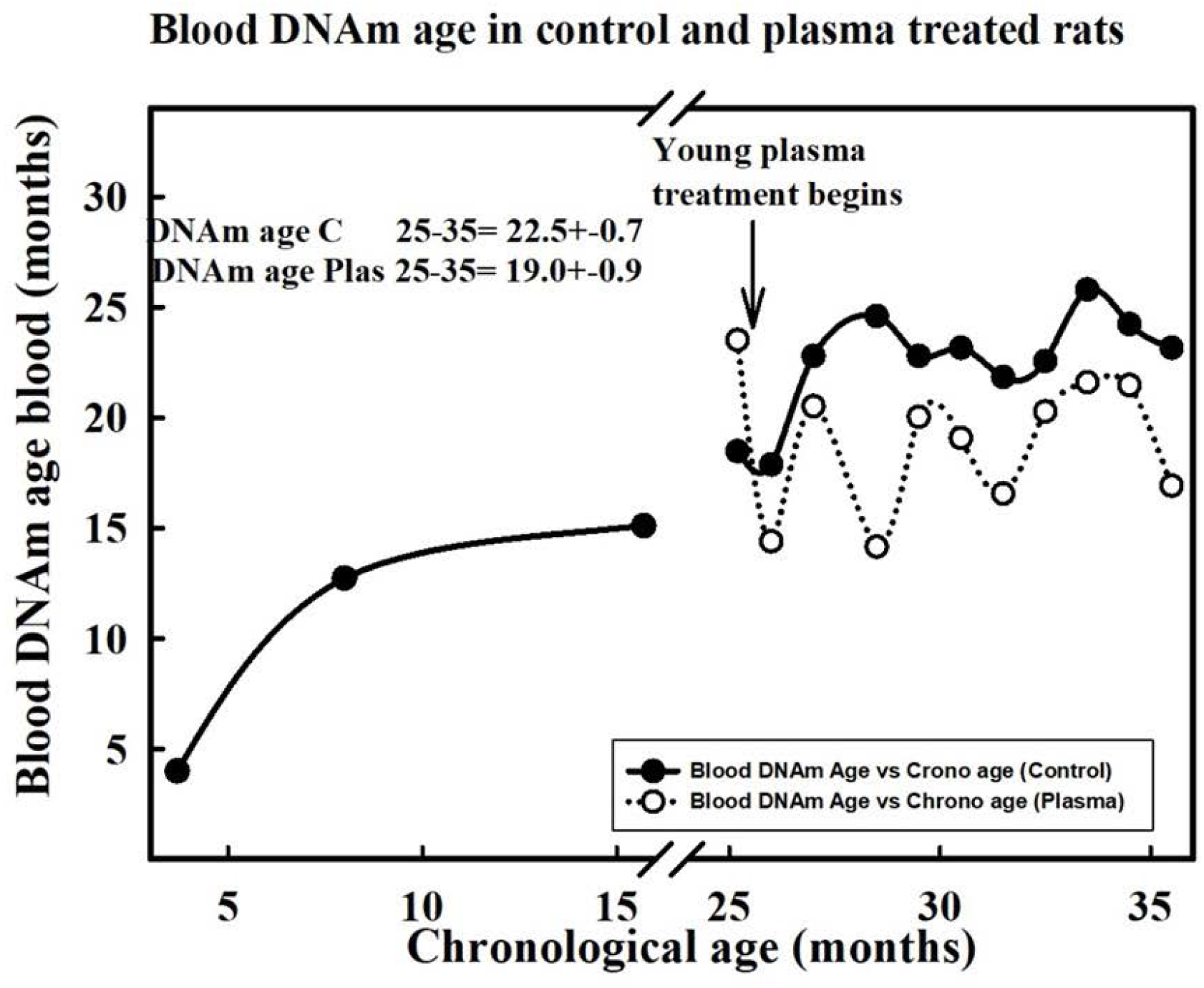
Progression of blood DNAm age versus chronological age in female rats treated with young plasma. The solid symbols correspond to untreated animals whereas the open circles represent DNAm age levels during the treatment. The data ranging from 3.7 to 15 chronological months correspond to a separate group of rats used to describe the life long evolution of blood DNAm age in intact females. The arrow indicates the time point where plasma treatment was started (25.6 months). Notice that from 27 to 35 months, DNAm age values increase much slower than chronological age in both control and treated animals.

### Effects of young plasma treatment on survival

In old rats (25 months) intraperitoneal injection of 1 ml young plasma every other week increased their average lifespan by 2.2 months as compared with untreated counterparts (**Fig. 2 upper plot).** From age 26 months to age 30 months no treated rats died whereas the survival curve of controls shows a steady decline from 26 to 30 months, a time period after which only one control rat remained alive. The surviving control was the longest-lived animal in the experiment, dying at age 35.8 months. Interestingly, the external appearance of the treated rats was healthier (their fur was whiter and shinier than that of controls. Also, treated rats were more active) than that of controls, the difference being more remarkable at the plateau of the survival curve **(Fig. 2, Photos 1and 2).**

**Figure 2.**
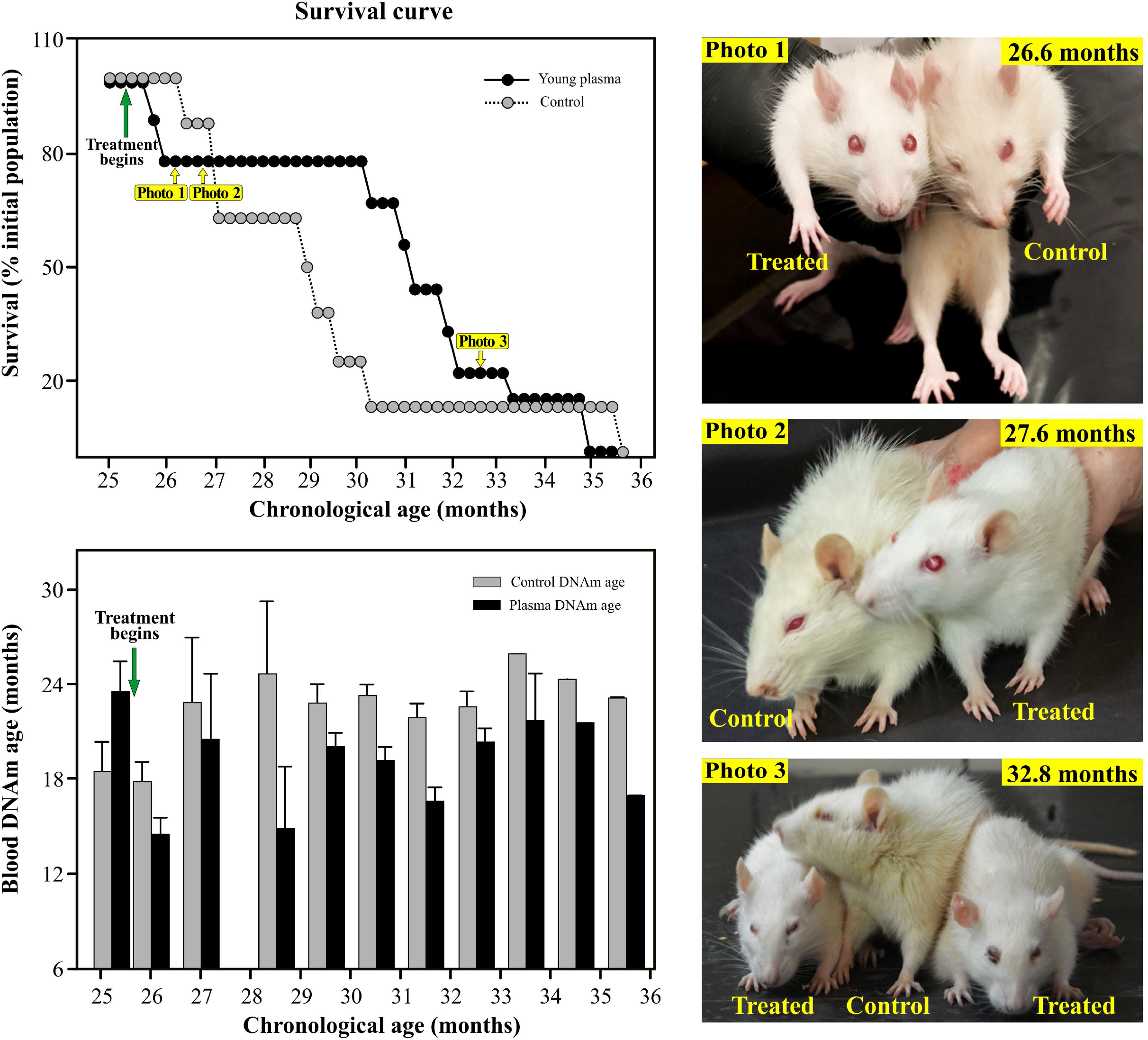
**Survival and epigenetic age of treated old rats-The upper graph** shows the survival curves of control and young plasma-treated rats. Yellow-labeled arrows indicate the time points at which rats were photographed. Notice that at age 26.6 and 27.6 months treated animals look younger than controls. **The lower plot** shows the DNAm age of blood samples taken at the indicated ages. After the beginning of the plasma treatment DNAm age became consistently lower in the treated animals.

### Effects of young plasma treatment on epigenetic age

Immediately after the start of young plasma administration, the DNAm age of the treated rats fell below the DNAm age of controls and remained consistently lower until the end of their lives **(Fig. 2, lower plot).** Although epigenetic age was consistently lower at all time points assessed, significant differences between control and treated groups were not detected in any pair of bars. On the contrary, comparison of average DNAm age in an age range spanning from 27 to 31.5 months and an age range spanning ages 32.5 to 35.5 months revealed that the former but not the latter, has a significant difference between control and treated groups **(Suppl. Fig. 1, lower plot).**

#### Differentially methylated CpGs

Assessment of each experimental group showed that the blood DNA methylation levels of 1638 CpGs were different between treated and control blood samples (q-value<0.05). **(Fig. 3 Panel A).** Of these, 1007 CpGs exhibited increased methylation (i.e., higher beta values), with age, while 631 CpGs showed decreased methylation levels **(Suppl. Tables 1 and 2, respectively).** Considering that the chip used here has approximately 106,000 CpGs on rat DNA sequences, the plasma treatement induced DNA methylation modifications in 1.6% of all rat CpGs present. Within the hypermethylated CpGs, 230 were located at promoter regions whereas within the hypomethylated CpGs, 155 were located at promoters regions **(Suppl. Tables 3 and 4, respectively)**.

**Figure 3.**
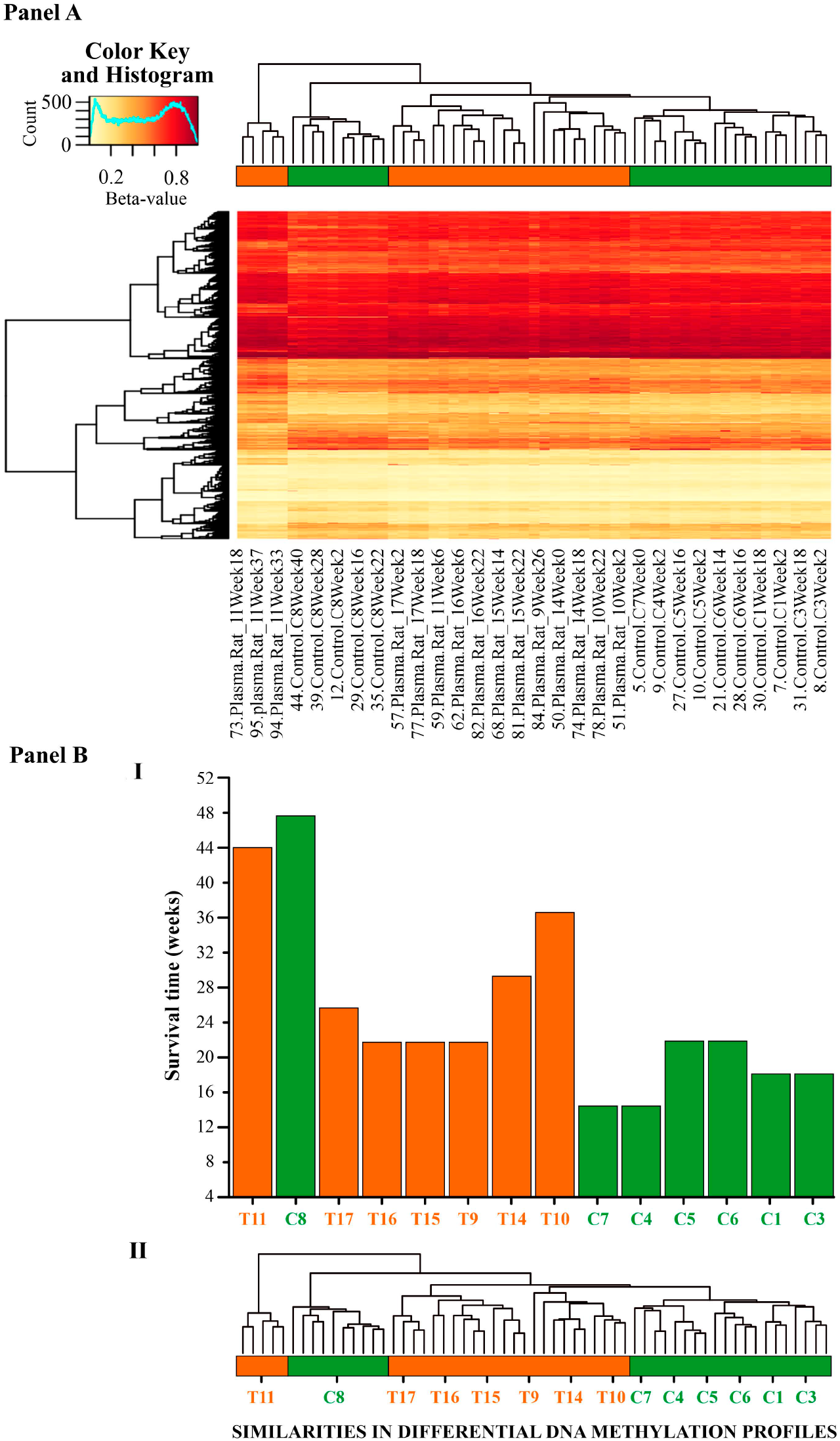
**Panel A (top).** Heatmap of differentially methylated CpG sites in the blood samples. Yellow denotes CpGs with the lowest methylation levels (beta values close to 0) and red denotes CpGs with the highest methylation levels (beta values close to 1). **Panel B (bottom)-** Barplot represents survival time in control and treated samples, displayed based on their similarity in the differential DNA methylation profile **(Panel B II).** In both the horizontal bar and barplot, color green corresponds to control rats whereas orange corresponds to treated animals. Notice that treated rats have similar differential DNA methylation profiles and are grouped on the left side of the plot. Similarly, control animals share similar differential DNA methylation profiles and group on the right side of the plot. Interstingly, control rat C8 is the longest lived animal and its differential DNA methylation profile shows a higher similarity with treated animals than with control ones. In terms of longer life and DNA methylation profile, rat C8 resembles more to treated than control rats.

We identified the CpGs in context with genomic feature: Promoter, Exon, Intron, Intergenic, 5’UTR, or 3’ UTR. The distribution profile of the CpG features in the genome is shown in **Fig. 4**.

**Figure 4.**
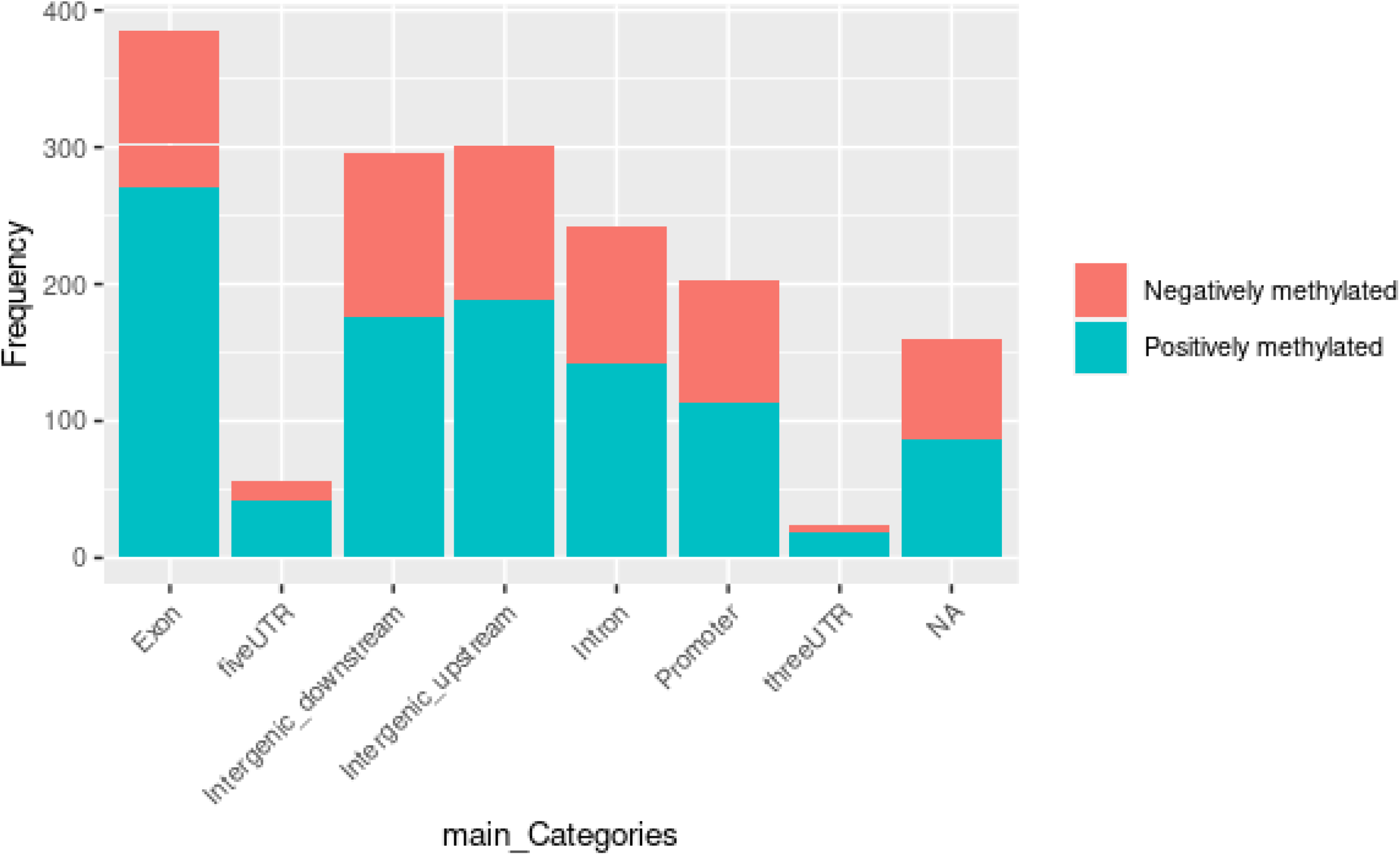
Genomic feature positions of the differentially methylated CpGs. Merged barchart of the genomic locations for the treatment-dependent positively methylated CpGs (pink color) and the negatively methylated CpGs (blue color).

In order to identify potential relationships between treatment-related differences in DNA methylation in rat blood, we performed a gene ontology (GO) enrichment pathway analysis. It showed no gene sets significantly enriched in differentially methylated CpGs. Interestingly a feature enrichment analysis of differentially methylated DNA sequences at promoter regions in genes involved in systemic regulatory activities revealed specific GO term enrichment related to insulin-like growth factors (IGFs) and to cytokines and chemokines associated with immune and homeostatic functions. The enriched sequences for IGFs were related to the IGF2 gene, the insulin receptor susbstrate (IRS1) involved in lifespan modulation in Drosophila and mice. S, an enriched gene related to IRS1, that may play a role in cell-cell adhesion processes was hypomethylated at the promoter region. There was also enrichment for the gene for insulin receptor susbtrate 2 (IRS2). The enriched sequences for cytokines and chemokines were CCL27 and CCL19, two genes encoding CC cytokine genes clustered on the p-arm of human chromosome 9. They elicit chemotactic effects by binding to the chemokine receptor CCR10 **(16).** They may play a role not only in inflammatory and immunological responses but also in normal lymphocyte recirculation and homing. They also may play an important role in trafficking of T-cells in thymus, and T-cell and B-cell migration to secondary lymphoid organs **(17)**. The gene SOCS2, encoding for a member of the suppressor of cytokine signaling (SOCS) family was also enriched. SOCS family members are cytokine-inducible negative regulators. SHC3, another enriched sequence codes for a gene related to the chemokine signaling pathway. There was enrichment in cytokine-cytokine receptor interaction CNTFR and INHBE, two genes related to the activity of cytokine genes CCL19, CCL21 and CCL27. SLAMF6 and ZBTB7B, two genes related with the regulation of interleukin-17 production were also enriched.

In general, differentially hypomethylated promoters were related with IGF-related gene promoters, whereas differentially hypermethylated promoters were related with chemo and cytokine gene promoters **(Suppl. table 5).**

### Similarities in DNA methylation profile within experimental groups

As expected, when rats were grouped according to the similarities in their differential blood DNA methylation profile through hierarchical clustering, samples from the treated and control rats clustered in separate groups**(Fig. 3, Panel B)**. On average, the treated rat samples corresponded to the longer lived animals wheras the control samples corresponded, on average, to shorter lived rats. An exception were the blood samples of control rat #8 which clustered with the treated samples. Therefore, in terms of DNA methylation profile, the C8 rat can be classified as a “treated-like control”.

## DISCUSSION

The present data support the hypothesis that in rats, young plasma conveys revitalizing factors that are able to reduce the epigenetic age of old animals which in turn may be causally related to an increased average lifespan and healthspan. In biological terms, it seems logical to assume that any treatment that increases lifespan will also improve healthspan. An interesting observation is that in young animals epigenetic time moves faster than physical time whereas in very old rats the ticking rate of the clock either markedly slows down or stops altogether. This is in line with reported observations in the rat hippocampus **(18)**. An immediate implication of this observation is that, if the clock does not tick in the old rats, the recorded decrease in epigenetic age induced by young plasma must be effected by forcing the clock to tick backward until reaching a lower level of DNAm age. Thus, the treated rats would be undergoing a significant, albeit modest, epigenetic rejuvenation. This is in line with a previous study reporting that treatment of old rats with a porcine plasma fraction, markedly setback their epigenetic age **(9).**

Incidentally, the observation that the epigenetic clock seems to stop ticking at advanced ages has a bearing on the hypothesis that the clock may be quasi-programming the aging process in mammals **(19).** In effect, in the context of this hypothesis, when the epigenetic clock of a senile mammal stops ticking, the developmental program runs out, and the ensuing deterioration of the homeostatic networks would be consequential to an epigenetic drift of the senile organism. The epigenome itself does not seem to deteriorate during the drift stage in very old mammals as suggested by the fact that somatic cells from old animals or even centenarians respond to rejuvenation genes and undergo full rejuvenation back to age zero, that is, they become induced pluripotent stem cells **(20–25)**. The idea that mammalian senescence may start when the developmental program runs out has been previously discussed **(26)**.

The present data suggest that the life-extending factors present in young plasma, whose concentration may drop as animals age, induce a significant but modest revitalization of the organism. Althogh this is a significant finding, there is documented evidence suggesting that adult mesenchymal stem cells (MSC) conveyed by young blood may possess a more potent life-extending capacity than young plasma. Thus, in the rat, bone marrow-derived human MSC were able to extend lifespan until 44 months **(27, 28)**, 8 months longer than the average maximum lifespan of albino rats **(29).** In F344 male rats it was reported that long-term intravenous treatment (one injection a month) of the animals with human amniotic membrane-derived mesenchymal stem cells (AMMSCs) or adipose tissue-derived mesenchymal stem cells (ADMSCs) extended their maximum life span from 23 months (saline-injected controls) to 30 and 34 months, respectively **(30)**.

In mice, it has been reported that bone marrow-derived MSC from young, but not old, mice implanted in full body X-irradiated old (18-24 mo.) female mice, moderately but significantly increased their lifespan and slowed their age-related loss of bone density **(31)**.

Another promising strategy for improving healthspan in elderly individiduals seems to be treatment with human umbilical cord plasma concentrates (HUCPC). Thus, in a phase 1 trial, it has been recently reported that intramuscular administration of HUCPC for 10 weeks to a group of 18 elderly subjects (average age, 74) improved the levels of different biomarkers, including epigenetic age measures, a number of clinical biomarkers of organ dysfunction, mitochondrial DNA copy number and leukocyte telomere length **(32).** The study also reports that umbilical cord blood plasma rejuvenates the kidney and leads to moderate epigenetic rejuvenation in human blood.

It has been reported that plasma dilution improves cognition and attenuates neuroinflammation in old mice, which suggests that dilution of putative circulating deleterious factors in old mice may be beneficial **(33)**, although there is no documented evidence that plasma dilution increases lifespan in mice or any other mammals. It seems unlikely that the blood dilution that took place in our rats after i.p. plasma injection every other week, may have played a role in the changes reported here. Besides, after 30 months of age, the treated rats began to die in spite of the fact that we continued injecting plasma until their natural death.

The control rat C8 was the longest-lived of both control and treated animals. It lived much longer than the second longest-lived control. We could draw an analogy from this rat with centenarians. Interestingly, the blood DNA methylation profile of this animal clustered with the blood samples from the treated rats **(Fig. 3 lower panel).** This observation shows that the blood DNA methylation profile of C8 was naturally similar to the profiles of the treated rats. In centenarians and their relatives, it has been shown that their epigenetic clock ticks slower than the clock of noncentenarians **(34)**. It seems therefore possible, that the epigenetic clock of C8 ticked slower than the clock of the other controls during its entire life.

It is also of interest that the biological effects induced in old rats by young plasma seem to have resulted from DNA methylation modification of only 1.6% of the total rat CpGs on the chip. This small fraction of differentially methylated CpGs may affect cells able to act at systemic level, an assumption in line with our results showing that young plasma-induced differential DNA methylation at promoters occurs in a number of IGF, cytokine and chemokine-related genes **(Suppl. Table 5)**.

It has been demonstrated in invertebrate species that the evolutionarily conserved insulin and IGF signaling (IIS) pathway plays a major role in the control of longevity. In mammals, different receptors exist that bind insulin, IGF-1 and IGF-2 with different affinities. Mutations that are associated with decreased growth hormone (GH)/IGF-1 signaling or decreased insulin signaling have been associated with enhanced lifespan **(35).** The GH/IGF-1 axis exerts an important modulatory effect on mammalian lifespan **(36).** For instance, Growth Hormone Receptor KO mice and Snell mice exhibit very Low epigenetic age compared to littermate controls **(36 bis)**.

IGF-1 is also a neuroprotective hormone able to rescue different dysfunctional neuronal populations in the brain of aging rats **(37, 38, 39).** Concerning the role of cytokines and chemokines in aging and longevity, many of these molecules regulate inflammation and other aspects of immune function **(40).** Cytokines act at systemic levels on multiple organs, including the brain. Thus, it has been reported that age-related changes in immune and neuroimmune system signaling impacts cognition **(41).** Furthermore, a recent study on heterochronic parabiosis in mice, identified hematopoietic stem and progenitor cells (HSPCs) as one of the most responsive cell types to young blood exposure, from which a continuum of cell state changes across the hematopoietic and immune system emanated, through the restoration of a youthful transcriptional regulatory program and cytokine-mediated cell-cell communications in HSPCs **(42).**

### Concluding Remarks

There is growing evidence that young plasma has a restorative effect in old rats **(43)** as well as in age-associated pathologies, particularly in the brain **(7, 44, 45, 46).** Thus, it has been reported that treatment with human cord plasma increases synaptic plasticity and hippocampal dependent cognition in aged mice and that one of the effectors of these improvements may be a circulating factor known as tissue inhibitor of metalloproteinases 2 (TIMP2) **(46)**. In old rats there is evidence that administration of plasma proteins may exert a marked reversal of epigenetic age **(9)**. In the present study we show, for the first time to our knowledge, that life-long treatment of old rats with young rat plasma induces a moderate but significant epigenetic rejuvenation, extends their mean lifespan and improves the healthspan of the animals, an effect that might be at least in part mediated by the IGF-1/IIS system as well as homeostatic cytokine and chemokine networks. We conclude that young plasma therapy may constitute a natural, minimally invasive intervention for epigenetic rejuvenation and health enhancement, readily translatable to the clinic.

## Supporting information

https://drive.google.com/drive/folders/1Xn0AcFOoVZgroxFQCpfgMYCUZCtiBkEb?usp=sharing

## ACKNOWLEDGEMENTS

The authors thank Dr. Martin Abba, CINIBA, UNLP, Argentina for technical comments on methylation analysis. The authors are indebted to Mr. Mario R. Ramos for design of the figures, to Ms. Yolanda E. Sosa for technical and editorial assistance and to DMV. Araceli Bigres for excellent care of our rat colony.

## Author Contributions

RGG, GRM and SH conceptualized the study; PC

## Data availability

The data that support the findings of this study are available from the corresponding author upon reasonable request.

## LEGENDS TO FIGURES

**Supplemental Figure 1.**
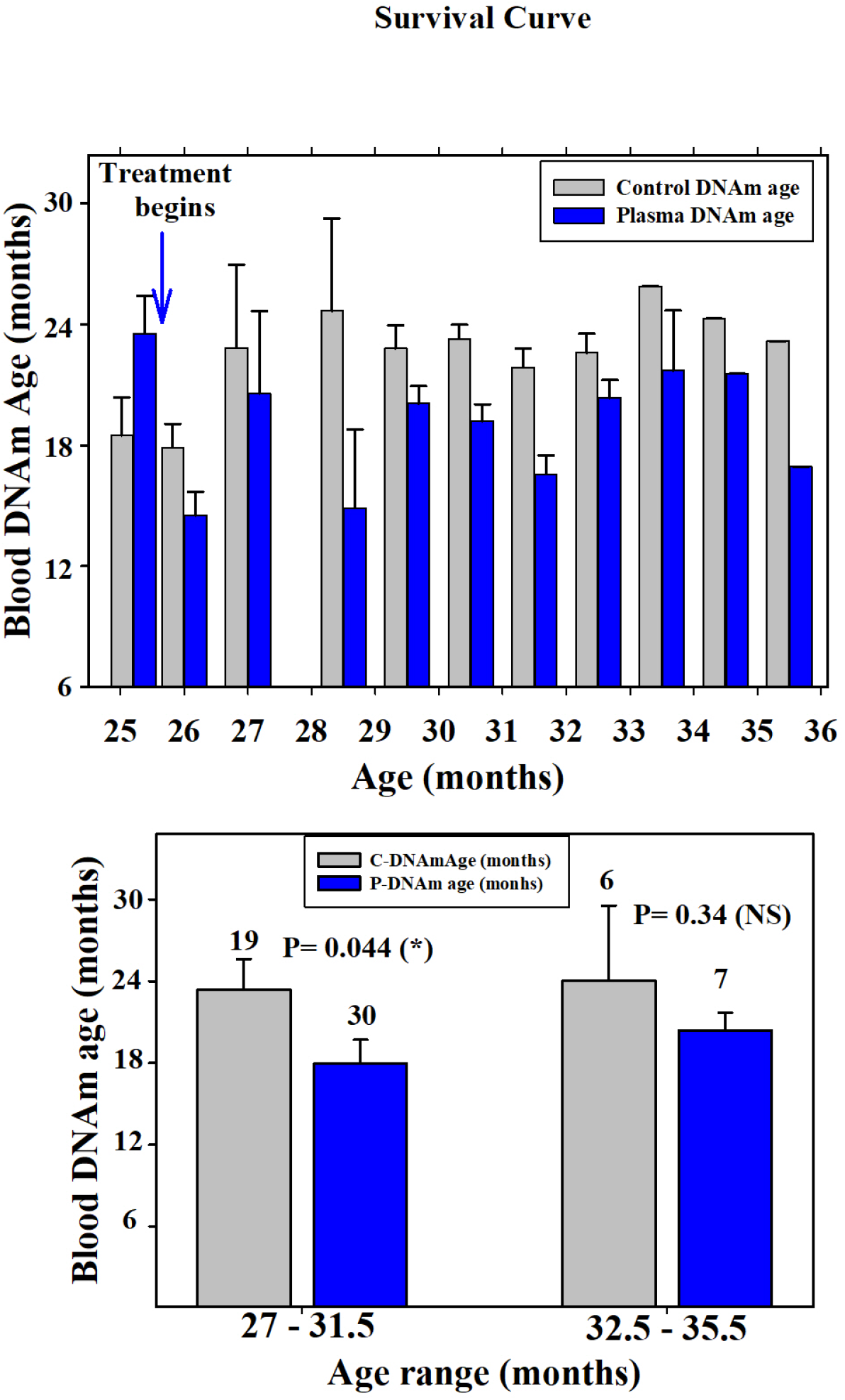
Statistical comparisons of epigenetic ages. The upper plot compares epigenetic age of control and treated rats at different time points of the study. None of the comparisons revealed significant differences. The lower plot assesses DNAm age in an age range spanning from 27-31.5 months versus an age range spanning ages 32.5-35.5 months. The former but not the latter shows a significant difference between control and treated groups.

**Supplemental Table 1-** Gene annotation of positively differentially methylated probes (qval<0.05).

**Supplemental Table 2-** Gene annotation of negatively differentially methylated probes (qval<0.05).

**Supplemental Table 3-** Gene annotation of positively differentially methylated CpGs present in promoters (qval<0.05).

**Supplemental Table 4-** Gene annotation of negatively differentially methylated CpGs present in promoters (qval<0.05).

**Supplemental Table 5-** Functional enrichment of differentially methylated genes in promoter regions of blood DNA from control and treated rats.

